# Snapshots of a shrinking partner: Genome reduction in *Serratia symbiotica*

**DOI:** 10.1101/057653

**Authors:** Alejandro Manzano-Marín, Amparo Latorre

**Affiliations:** Institut Cavanilles de Biodiversitat I Biologia Evolutiva – Universitat de València, Genética Evolutiva, Paterna, 46980, Spain; Fundación para el Fomento de la Investigación Sanitaria y Biomédica de la Communitat Valenciana (FISABIO), Genómica y Salud, València, 46020, Spain

## Abstract

Genome reduction is pervasive among maternally-inherited endosymbiotic organisms, from bacteriocyte- to gut-associated ones. This genome erosion is a step-wise process in which once free-living organisms evolve to become obligate associates, thereby losing non-essential or redundant genes/functions. *Serratia symbiotica* (Gammaproteobacteria), a secondary endosymbiont present in many aphids (Hemiptera: Aphididae), displays various characteristics that make it a good model organism for studying genome reduction. While some strains are of facultative nature, others have established co-obligate associations with their respective aphid host and its primary endosymbiont (*Buchnera*). Furthermore, the different strains hold genomes of contrasting sizes and features, and have strikingly disparate cell shapes, sizes, and tissue tropism. Finally, genomes from closely related free-living *Serratia marcescens* are also available. In this study, we describe in detail the genome reduction process (from free-living to reduced obligate endosymbiont) undergone by *S. symbiotica*, and relate it to the stages of integration to the symbiotic system the different strains find themselves in. We establish that the genome reduction patterns observed in *S. symbiotica* follow those from other dwindling genomes, thus proving to be a good model for the study of the genome reduction process within a single bacterial taxon evolving in a similar biological niche (aphid-*Buchnera*).

## Introduction

Obligate microbial symbionts (weather primary, secondary, tertiary, or other) are present in a variety of eukaryotic organisms, such as leeches (Annelida: Hirudinida)^1^, gutless oligochaetes (Annelida: Oligochaeta)^2^, and insects (Arthropoda: Insecta)^3^ (see^4^). These have the capacity to produce essential nutrients their hosts cannot synthesise nor obtain from their diet^5–8^, making them essential for the correct development and survival of their partners. On the other hand, facultative symbionts are dispensable, although under certain environmental challenges/niches, they can endow the host with desirable traits, ranging from defence against parasitoids or fungal parasites to survival after heat stress (reviewed in^9, 10^). Moreover, these facultative endosymbionts can even affect the performance of its host on a certain food source (e.g. a plant)^11–13^.

Whichever the symbiont’s function, taxonomic position, or status (facultative or obligate), a common feature from maternally inherited endosymbiotic organisms is the possession of a reduced genome, when compared to their free-living counterparts^8, 14–22^. The sequencing of these genomes has undoubtedly provided important clues into the distinct features these display along the erosion process. While mildly-reduced genomes such as the one from *Sodalis glossinidius*, (facultative endosymbiont of the tsetse fly *Glossina morsitans morsitans*), *Sodalis pierantonius* (primary obligate endosymbiont from the rice weevil *Sitohpilus oryzae*), and *Hamiltonella defensa* (facultative endosymbiont from the aphid *Acyrthosiphon pisum*) show intermediate guanine-cytosine (hereafter **GC**) contents and a massive presence of both mobile elements (hereafter **MEs**) and pseudogenes^23–25^, highly reduced genomes such as the ones from *Buchnera* (primary obligate endosymbiont of aphids) and *Blochmannia* (primary obligate endosymbiont from carpenter ants) are highly compact with few pseudogenes and show no traces of MEs^15, 26^. However valuable the study of these genomes is, very few examples are available for different bacterial strains, belonging to a single genus, holding differentially-reduced genomes. These examples include *Arsenohponus* symbionts of a parasitic wasp (*Nasonia vitripennis*) (INSDC:AUCC00000000.1), the brown planthopper (*Nilaparvata lugens*)^27^, and louse flies (Diptera: Hippoboscidae)^28^ (INSDC:CP013920.1); *Coxiella* symbionts of ticks^21, 22^; and *Sodalis* symbionts from the tsetse fly^23^ and rice weevil^24^.

*Serratia symbiotica*, a secondary endosymbiont harboured by many aphids^29–32^, is particular among currently sequenced bacterial symbionts. While strains harboured by *Aphis fabae* and *Acyrthosiphon pisum* (Aphidinae subfamily) are of facultative nature^29, 33, 34^, strains from the aphids *Cinara tujafilina*, *Cinara cedri*, and *Tuberolachnus salignus* (Lachninae subfamily) have established co-obligate associations with both the aphids and its primary obligate endosymbiont, *Buchnera*^35–37^. This co-obligate association was putatively triggered by a loss of the riboflavin biosynthetic genes in *Buchnera* from the Lachninae last common ancestor^31^. In addition, depending on the strain, its cell shape is either rod-like (strain CWBI-2.3 from *Ap. fabae^38^*, hereafter **SAf**), filamentous (strain Tucson from *Ac. pisum^39^*, hereafter **SAp**; and strain SCt-VLC from *C. tujafilina^35^*, hereafter **SCt**), or spherical (strain SCc from *Cinara cedri^36^*, hereafter **SCc**; and strain STs-Pazieg from *T. salignus*^31^, hereafter **STs**). Not surprisingly, these strains hold genomes of contrasting sizes, ranging from 3.58 to 0.65 mega base pairs (hereafter **Mbp**)^16^,^20^,^35–37^. Furthermore, although most strains have not yet (to our knowledge) been cultured, similarly to many insect obligate endosymbionts (reviewed in^40^), SAf is able to grow freely in anaerobic conditions on a rich medium^38^. Finally, while the phylogenetic relations of *S. symbiotica* are not fully resolved^31^, they show a clear sister relationship to *Serratia marcescens*, a species comprised of various free-living bacterial strains for which complete genomes are available.

In the current study, we analysed the genomes of currently-available *S. symbiotica* strains and compared them to the free-living insect pathogen *S. marcescens* strain Db11 (hereafter **Db11**)^41^. Through comparative genomics we investigated genome rearrangement, the enrichment, and loss, of MEs, and the erosion undergone by RNA features and the informational machinery in *S. symbiotica*. Additionally, we describe the diminution of certain genes and the possible functional consequences of these reductions. Finally, we relate all these features to different stages of the symbionts’ integration to the aphid-*Buchnera* symbiotic consortia and discuss the features which are convergent with other dwindling endosymbiotic genomes.

## Results and Discussion

### *S. symbiotica* strains and their shrinking genomes

Generally, “ancient” obligate endosymbionts hold highly reduced genomes as small as 112 kilo base pairs^42^ (hereafter **kbp**), conversely, more “recently” derived endosymbionts (including facultative ones) tend to display larger genomes, all the way up to the 4.5 Mbp genome of *S. glossinidius* (reviewed in^43^). Accordingly, the different genomes of *S. symbiotica* strains land within and along this spectrum, from the large 3.58 Mbp genome of the facultative SAf to the small 0.65 Mbp genome of the co-obligate STs (Figure 1). Similarly to the other large endosymbiotic genomes^23–25, 44, 45^, SAf, SAp, and SCt’s display a large enrichment of MEs, both in terms of diversity and number of them. Interestingly, the composition of the insertion sequence (hereafter **IS**) families (the most common type of MEs found within these genomes) seems to be lineage-specific. While IS3 and IS256 are the most prevalent in SAf and SAp (both facultative endosymbionts from Aphidinae aphids), IS481 and IS5 are the most common in SCt (co-obligate endosymbiont from *C*. tujafilina [Lachninae]). Conversely, the smaller genomes of SCc and STs lack any traces of MEs, congruent with similar-sized endosymbiotic genomes (see^3^). Following the trend of many other dwindling genomes^3^, all *S. symbiotica* have a GC content lower than that of their free-living counterpart, Db11. While this GC content is very similar among SAf, SAp, and SCt (52.1%), there is a marked decrease in SCc (29.2%) and even more so in STs (20.9%). Additionally, while there is a great enrichment of pseudogenes in SAf (126), SAp (550), SCt (916), and SCc (110), the small STs is almost deprived of these gene remnants. This genetic erosion comes along with a decrease in coding density. Accordingly, while SAf shows only a small decrease when compared to Db11 (87.9% to 78.2%), SAp, SCt, and SCc exhibit a marked drop down (56.8%, 53,4%, and 39.0%, respectively), mainly due to the increased pseudogenisation and “junk” DNA. On the other hand, the highly-reduced STs shows a high coding density (77.5%). This difference between SCc and STs is mainly due to high-amount “junk” DNA that is present in SCc’s genome, amounting to almost half of it^36^. Finally, we also found a gradual loss (from free-living Db11 to co-obligate intracellular STs) of RNA features (rRNAs, tRNAs, and other non-coding RNAs [hereafter **ncRNAs**]), revealing their different levels of genomic erosion. As has been previously observed in other endosymbionts, genome erosion comes with a “disturbance” of the functional profile of the organism, when compared to their free-living relatives^14, 18^. Accordingly, prior analyses have described that while the functional profiles of free-living *Serratia* strains were very stable, a displacement of it was evident in SCc and SAp^46^. Through a similar analysis using all five currently available *S. symbiotica* strains, we have found that while the recently-derived SAf, SAp, and SCt strains cluster together forming a sister group to Db11, the highly-reduced SCc and STs form a divergent cluster from the rest of *Serratia* strains (Figure 2A). These two *S. symbiotica* clusters differ mainly in the relative presence of MEs (category **X**) and translation-related genes (category **J**). While the former reflects the enrichment of SAf, SAp, and SCt’s in MEs, the latter evidences the common trend in highly reduced endosymbionts to retain housekeeping genes (e.g. category J includes all ribosomal proteins) (see^3^).

**Figure 1.**
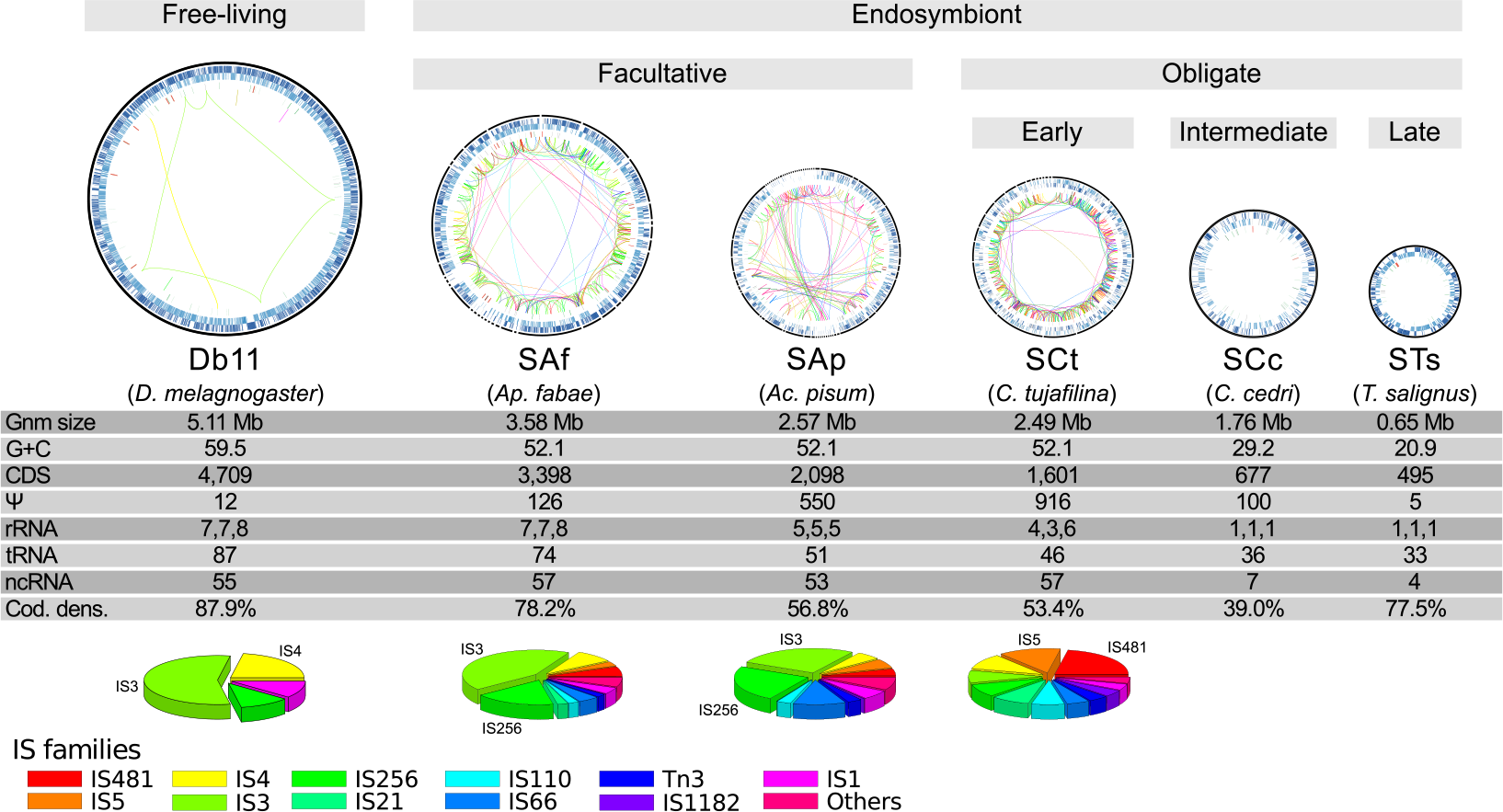
Genome reduction in *S. symbiotica.* *Serratia* genomes are depicted as circular plots and are arranged from largest (leftmost) to smallest (rightmost). From outermost to innermost, the rings within the genome plots display features on the direct strand, the reverse one, and RNA genes. Inside the circles, coloured lines connect the same-family IS elements scattered throughout the genome, following the colour code at the very bottom of the image. The grey bars on top of the genome plots describe the lifestyle and genome reduction stage. Underneath the genome plots, the strain alias and the host, between parenthesis, are shown. Below, a table showing the genomic features of each strain and pie charts displaying the relative abundance of IS-family elements, with the two most abundant highlighted by name. Underneath, the colour code for the different IS elements.

**Figure 2.**
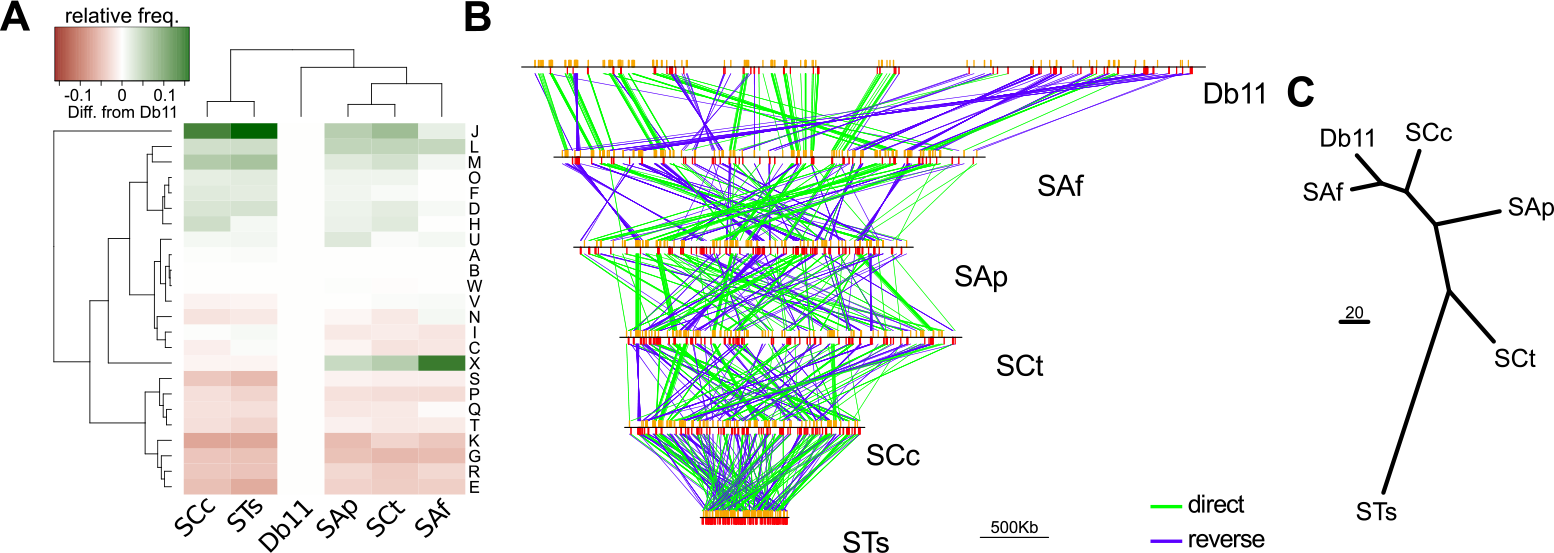
*S. symbiotica* functional profile displacement and genome rearrangement. (**A**). Heat map showing the two-way clustering of the *S. symbiotica* COG profile’s differences, relative to the free-living *S. symbiotica.* On the right, one-letter code for the COG categories. (**B**) Graphic linear representation of the rearrangements undergone in *S. symbiotica.* Orange and red vertical bars mark the position of single-copy conserved genes. Contigs that do not have single-copy conserved genes are not displayed. (**C**) Unrooted tree as calculated by MGR for the minimum number of rearrangements undergone by *S. symbiotica*.

In the early stages of an endosymbiont’s genomic reduction, the genome’s enrichment in MEs can lead to rearrangement^18, 35^. These rearrangements get fixed in the endosymbiotic lineage once the MEs have been lost, as is observed by the general genome-wide synteny displayed in *Buchnera*^15, 47^, *Blochmannia*^26^, or *Blattabacterium*^48^. Free-living *Serratia* strains display general genome-wide synteny^46^, on the contrary, *S. symbiotica* genomes display various rearrangements when compared to free-living Db11’s, and even among each other’s (Figure 2B and C). Interestingly, while the less-reduced genome of SAf displays the closest relationship (in terms of rearrangements) to Db11, the drastically-reduced genome of STs has accumulated the highest number of rearrangements. Also, SCc and STs’ genomes, which both lack MEs, display no synteny between them. These observations suggest that all *S. symbiotica* lineages have diverged before the loss of MEs, allowing a great number of lineage-specific reorganisation.

### Erosion of essential amino acid biosynthetic routes

A general feature of endosymbiotic genomes is the loss of non-essential genes, leading to highly reduced genomes with a genetic repertoire specialised in the symbiotic function (reviewed in^3^). In aphids, *Buchnera*, the primary obligate endosymibont, is mainly in charge of producing essential amino acids (hereafter **EAAs**) for its host. Therefore, it is expected that co-existing symbionts show degraded biosynthetic routes involved in the production of these compounds. By analysing these routes in *S. symbiotica*, the gradual degradation of genes and operon attenuators implicated in the synthesis of EAAs becomes immediately evident (Figure 3). The recently-derived SAf shows intact routes for most EAAs, with the notable expections of lysine and methionine. As already described in a previous study, there is a marked difference in the retention of leucine, arginine, and histidine biosynthetic-related genes even between the closely related facultative SAp and co-obligate SCt^35^. Finally, by comparing SCc and STs against the other *S. symbiotica* strains and each other, it becomes evident that both have become highly dependent on *Buchnera* for the supply of EAAs, with the main difference between SCc and STs being the purging of the remaining pseudogenes in the latter.

**Figure 3.**
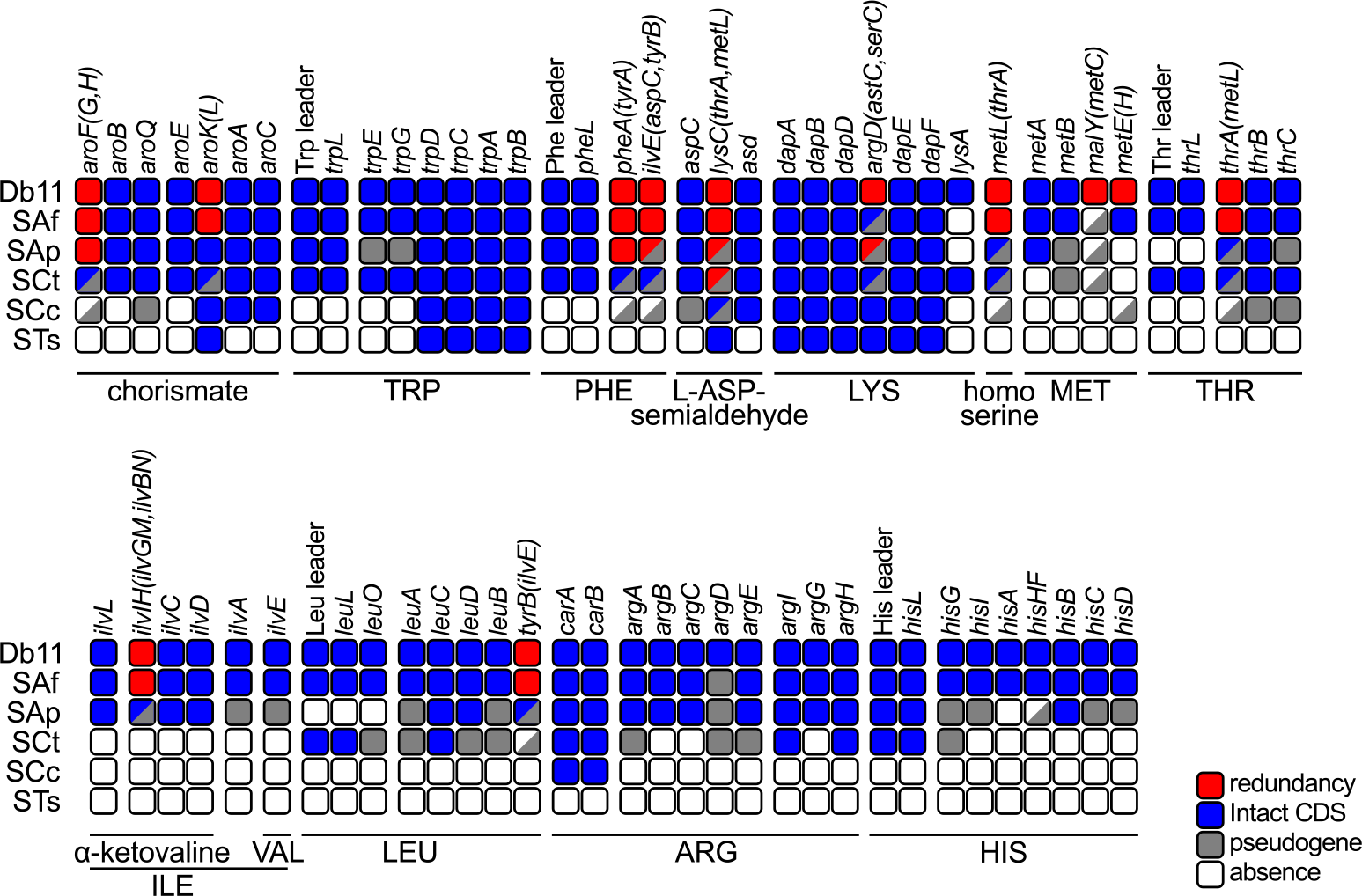
Erosion of essential amino acid biosynthetic genes in *S. symbiotica.* Inactivation tables showing the genes and leader sequences involved in the essential amino acid biosynthetic routes in *S. symbiotica* genomes compared to free-living Db11. At the top and left of the table, gene names for each enzymatic step and abbreviation for each *Serratia* strain, respectively. At the bottom of the table, black lines encompass the enzymatic steps required for the biosynthesis of each compound. Amino acid names are displayed using standard three-letter abbreviations. At the bottom-right, colour code for squares. Half-coloured boxes mean the genes catalysing the enzymatic step are present in different states.

### Decay of RNA features and the loss of regulation

Typically, highly-reduced endosymbionts retain only a small number of ncRNAs and other RNA features (see Supplementary **Figs. S1** and **S2**). Through an annotation of these in the genomes of *S. symbiotica* and Db11, we have explored the erosion of RNA features (Figure 4: top panel). We found that in the recently-derived endosymbionts SAf, SAp, and SCt, many of these features are still retained, although differentially. This points towards drift acting behind the loss of these features at the early stages of genome reduction. As expected, these three genomes show the acquisition of ME related ncRNAs, which all belong to the large class of self-catalytic group II introns (RF00029, RF01999, RF02001, RF02003, RF02005, RF02012). In the intermediate and advanced stages of drastic genome reduction SCc and STs find themselves in, respectively, most of the RNA features have been lost. Conserved features across *S. symbiotica* are the the 4.5S RNA component of the signal recognition particle (SRP) (*ffs*), the RNase PM1 RNA component (*rnpB*), the tmRNA (*ssrA*), the tpke11 small RNA (of unknown function), the leader sequence from the *rnc-era* transcription unit (coding for the ribonuclease 3 and the GTPase Era), and the alpha operon leader (coding for the 30S ribosomal subunits S13, S11, and S4; the 50S ribosomal subunit L17; and the DNA-directed RNA polymerase subunit alpha). The first three are interestingly also retained in other small genomes, but unidentifiable in some tiny genomes (Supplementary **Fig. S1**), hinting at these being essential functions retained until the last stages of genome reduction. Since most of these RNA features are related to the regulation of gene expression (small antisense RNAs, riboswitches, and leader sequences [including amino acid operon attenuators]), these losses would reflect a general trend of gene-regulation-loss in endosymbiotic genomes through the erosion of RNA features.

Regarding tRNAs, we observed a drastic reduction in tRNA-gene number, particularly marked in SCc and STs (Figure 4: bottom panel). These losses, as in other reduced endosymbionts (see Supplementary **Fig. S2**), mainly affect redundancy rather than variety. Contrasting the other *S. symbiotica* genomes, we were unable to detect a tRNA with aminoacyl charging potential for glutamate in SCc. This is similar to what is observed in other tiny genomes, where some tRNAs with certain aminoacyl charging potential are absent (Supplementary **Fig. S2**). However, the presence of a tRNA^Glu^ in a yet-unidentified plasmid cannot be discarded. Also, a loss of the selenocysteine tRNA is already present in the early co-obligate SCt, consistent with the loss of other selenocysteine-related genes, and completely absent in the smaller SCc and STs. It is important to remark that, in SAp, one of the tRNA^Met^ copies has undergone a mutation in its anticodon (<u>C</u>AT→<u>A</u>AT), which could theoretically lead to the ATT codon to be recognised as coding for methionine. Finally, the tRNAs for formyl-methionine (tRNA^fMet^), in charge of aminoacylation of the starting methionine, and lysylated isoleucine (tRNA^kIle^) are conserved even in the two smallest *S. symbiotica*. This follows the trend observed in other reduced genomes (Supplementary **Fig. S2**) and points towards the essential nature of these tRNAs.

**Figure 4.**
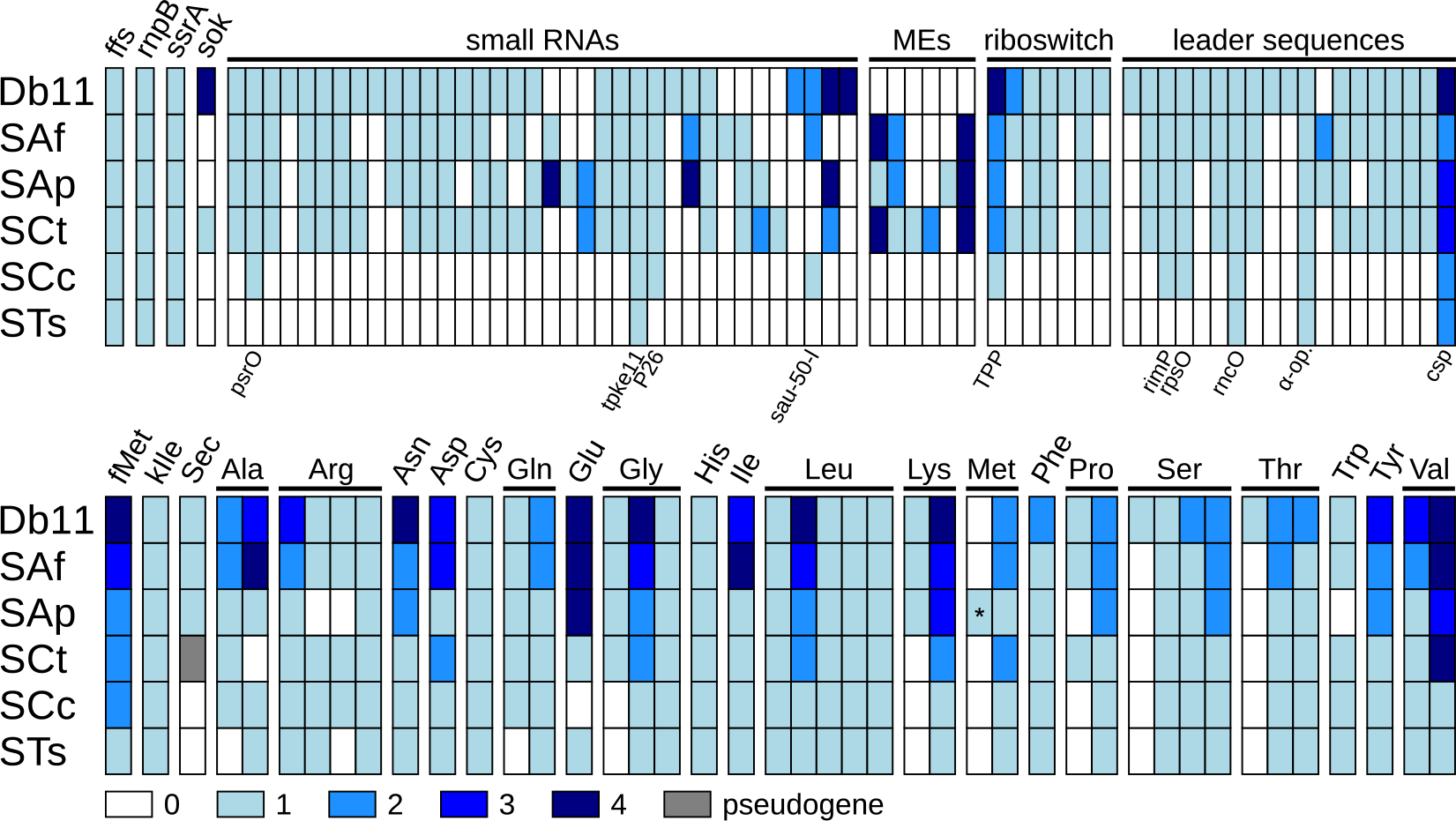
Decay of tRNA and other RNA features in *Serratia* genomes. (**Top**) Colour-coded diagram showing the decay of RNA features in different *S. symbiotica* genomes. On top of the matrix, gene names (for the first four columns) and RNA categories (for the rest) are indicated. On the bottom of the matrix, feature names are indicated for those features retained in SCc and STs. (**Bottom**) Colour-coded diagram showing the decay of tRNA features in different *S. symbiotica* genomes. On the top of the matrix, aminoacyl charging potential for each tRNA species (as inferred by TFAM). Each column represents a different anticodon. Asterisks indicate putative codon reassignments as judged by **TFAM**. fMet= *N*-Formylmethionine, kIle=lysylated isoleucine.

### Informational machinery

By analysing and comparing the informational machinery (ribosome-, transcription-, translation-, and DNA replication/repair-related genes) in *S. symbiotica* strains, both high preservation as well as gradual patterns of deterioration become evident in different categories. The ribosome, as well as the tRNA aminoacylation genes are mostly perfectly preserved (Figure 5: top). Marked differences include the presence of multiple copies of the three rRNA genes in SAf, SAp, and SCt, and the absence of two ribosomal proteins (*rpsI* and *rplM*) as well as the prolyl-tRNA synthetase gene (*proS*) in STs. While the retention of only one copy of the rRNA genes reflects the tendency of endosymbiotic, and other reduced genomes, to eliminate redundancy^49^, the loss of the *rpsI* gene (coding for the 30S ribosomal subunit S9) reflects the loss of a non-essential gene. In *Escherichia coli*, it has been experimentally proven that a null mutant of the *rplI* gene is able to grow, albeit showing a slow growth phenotype^50, 51^. Most intriguing are the losses of the *rplM* (coding for the 50S ribosomal subunit L13) and *proS* genes. The former has been described as essential in *E. coli*^51^, and its loss could be related to the loss of *rpsI*, that together with *rplM* forms an operon. The latter, could reflect a putative functional replacement of the ProS protein activity by another non-specific aminoacyl-tRNA synthetase. This phenomenon has been observed for the prolyl-tRNA synthetase of *Deinococcus radiodurans*, which has the ability to charge cysteine to tRNACys^52^. This non-specific aminoacyl-tRNA synthetases have also been observed in archeal organisms (reviewed in^53^), suggesting this to be a common mechanism to cope with the lack of a specific aminoacyl-tRNA synthetase.

**Figure 5.**
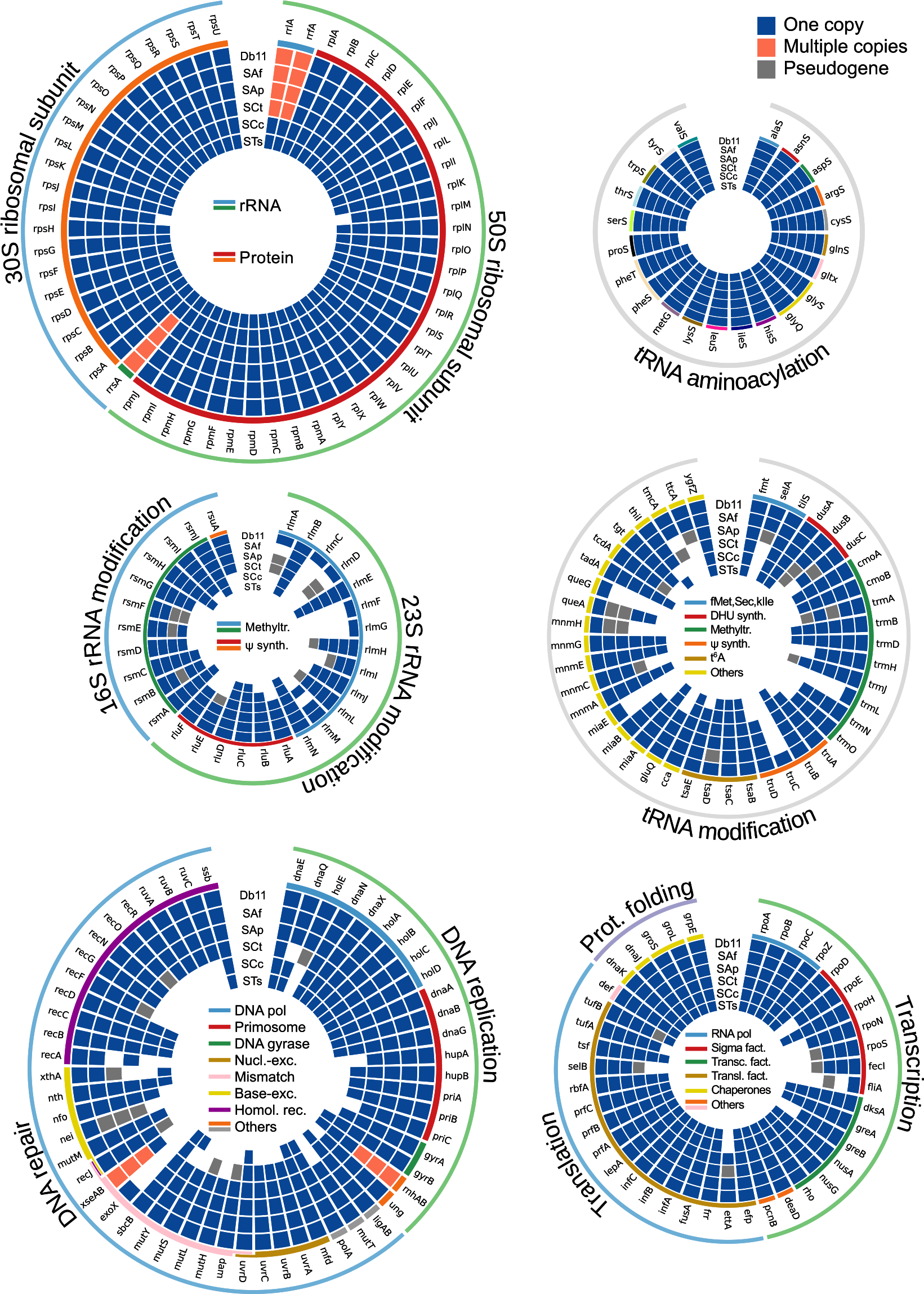
Informational machinery in *S. symbiotica* genomes. Circular plots displaying the different genes involved in functional categories (top left, ribosome; top right, tRNA aminoacylation; middle left, rRNA modifications; middle right, tRNA modification; bottom left, DNA replication and repair; bottom right, transcription and translation) in the informational machinery of *S. symbiotica* strains and Db11. Outer lines in each circular plot delimit the subcategory. Form outer to inner, the rings in the plot stand for the gene name, the colour-coded lines delimiting categories/complexes, and boxes standing for the presence or absence of the genes in Db11, SAf, SAp, SCt, SCc and STs.

Both rRNAs and tRNAs undergo a series of modifications that are required to produce the mature version of these ncRNAs (reviewed in^54, 55^). By analysing the genes involved in both rRNA and tRNA modifications, we observed that while the recently-derived SAf, SAp, and SCt hold a rather complete set (with particularly marked losses of 23S rRNA methyltransferases), the highly-reduced SCc and STs retain only a small fraction of these genes (Figure 5: middle). With the notable exceptions of the *fmt*, *tilS*, *trmD*, *tsaB*, *tsaC*, *tsaD*, and *tadA* genes (all retained in the small SCc and STs), individual knockout mutants all of the rRNA and tRNA modification-related genes in *E. coli* (except *miaE*, which is not present in this organism) have dimmed them non-essential^56–59^. The *fmt* and *tilS* genes code for the proteins responsible for the attachment of a formyl group to the free amino group of methionyl-tRNA^fMet^ (for initiator methionine)^60^ and the modification of the wobble base of the CAU anticodon of the tRNA^kIle61, 62^, respectively. The retention of these two genes thereby insure both the correct charging of the initiator methionine in proteins (which is posttranslationally-removed) and the accurate recognition of the AUA codon as coding for Isoleucine. On the other hand, the genes *tsaB*, *tsaC*, and *tsaD* (along with the *tsaE* gene) are responsible for the biosynthesis of the threonylcarbamoyladenosine (t6A) residue at position 37 of ANN-decoding tRNAs^63^. Interestignly, the *tsaE* gene (an ATPase), which has been found to be non-essential in *E. coli* under anaerobic conditions^64^, is missing from STs, thus the biosynthesis of t6A would be either putatively impaired or working in an unknown way. Finally, the *tadA* gene, which codes for a tRNA-specific adenosine deaminase that is essential for viability in *E. coli^65^*, is retained even in the small STs.

In regards to DNA replication and repair, the gene losses are particularly marked in the most genomically reduced symbionts, SCc and STS, affecting mostly DNA repair-related genes (Figure 5: bottom left). This is also observed in other reduced endosymbionts (see^43^), and is possibly related to the triggering of more drastic genome erosion (reviewed in^3^). DNA replication-related losses affect the non-essential *holE* gene of the DNA polymerase, the priA-dependent primosome (retaining an elementary DNA-dependent one [missing the auxiliary Hup proteins]), and the *gyrA* subunit of the DNA gyrase. These latter, although identified as essential in *E. coli^57^*, has also been found to be missing from tiny genomes^43^, thereby suggesting its function could be taken over by an alternative enzyme or it actually being non-essential in some endosymbiotic organisms.

In terms of transcription-and translation-related genes, a high-degree of retention in all *S. symbiotica* genomes can be observed (Figure 5: bottom right). Gene losses mainly affect the sigma factors, with STs retaining only the *rpoD* and *rpoH* genes, coding for σ^70^ and σ^32^, respectively. While the former is generally preserved in endosymbionts^43^, the latter is missing from endosymbionts such as *Blattabacterium* and *Nasuia*. σ^32^ is required for the normal expression of heat shock genes and for the heat shock response through the regulation of the synthesis of heat shock proteins^66^, and thus its retention/loss could be specific of certain endosymbiotic systems.

### Dwindling genes: stripping proteins down to the bones

Through the manual curation of the annotation of SCt, SCc, and STs endosymbionts^35, 37^, we noted that some genes (*atpC*, *cysJ*, *deaD*, *dnaX*, *ftsN*, *hscA*, *metG*, *pcnB*, *rnr*, and *tolC*) seemed to be shorter in STs, and sometimes consistently shrunken across *S. symbiotica*, compared to those of free-living *E. coli* and even Db11. However, while these genes showed truncated or missing domains, they displayed a high degree of sequence conservation, when compared to Db11. Thorough examination of these shrunken genes revealed that experimental evidence, mainly form *E. coli*, have proven that truncated versions of these proteins were able to function with few to none obvious phenotypic consequences (details recorded in the annotation files available from the INSDC). Particularly evident is the loss of non-essential domains in six proteins: AceF, DnaX, FtsK, FtsN, and Rnr (Figure 6). The AceF protein (E2 component of pyruvate dehydrogenase complex) has undergone the loss of one or two biotin/lypoyl domains (PF00364) in all *S. symbiotica*, namely STs retains only one. In *E. coli*, it has been shown, through the *in vitro* deletion of biotin/lipoyl domains, that one single domain suffices with respect to enzyme activity and protein function^67^. The tau subunit of the DNA polymerase III is coded by the *dnaX* gene, however an alternative isoform, denominated gamma subunit, is produced due to a programmed ribosomal frameshifting, which leads to a premature stop codon in the −1 frame at codon 430^68^. *in vitro* experiments with the shorter isoform, which lacks the tau 4 and 5 domains (PF12168 and PF12170), indicate that gamma is to be sufficient for replication^69^. The most drastic gene diminutions are observed in the *ftsK* and *ftsN* genes (whose products are involved in cell division), where SCc and STs preserve only a very small portion of the original gene. Independent *in vivo* experiments in *E. coli* mutants coding only for truncated FtsK (amino acids 1-200)^70^ or FtsN (amino acids 1-119)^71^ proteins, have corroborated that these tiny versions are sufficient for cell division, although short to long filamentous cells were observed to occur. Regarding MetG (Methionyl tRNA-synthetase), both SCc and STs are lacking the C-terminal putative tRNA binding domain (PF01588). Genetic complementation studies and characterization of C-terminally truncated enzymes in *E. coli*, established that MetG can be reduced to 547 residues without significant effect on either the activity or stability of the enzyme^72^. Finally, a deletion of the C-terminal basic domain of the Rnr protein (ribonuclease R) can be observed only in STs. This could lead to an increase in activity of this enzyme, since assays using purified truncated Rnr proteins from mutant *E. coli*, lacking the 83 residues from the C-terminus, were shown to display higher affinity and *circa* 2-fold higher activity than full length wild-type Rnr (on poly[A], A[17] and A[4] substrates)^73^. Through the alignment of the aforementioned putatively-functional proteins against other small and tiny genomes, we corroborated most of these gene diminutions are common among these organisms (Supplementary **Data S1**). This suggests that selection might favour gene diminution (the retention of only essential domains of a protein), relaxing selective constraints in non-essential gene regions, thus further contributing to genome reduction.

**Figure 6.**
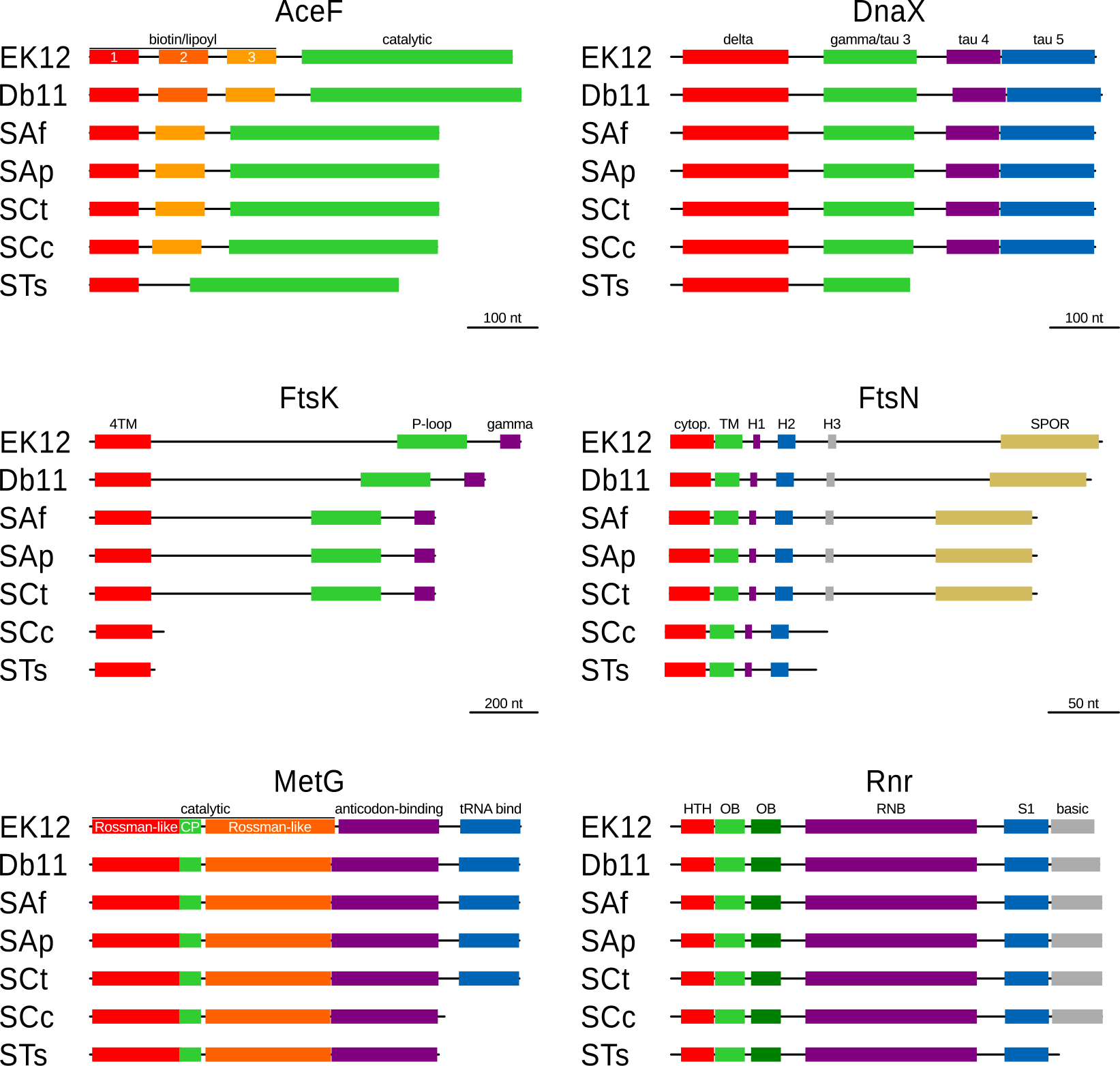
Diminution of genes in S. *symbiotica*. Graphic representation of the proteins that show evident loss of non-essential domains (as judged by experimental evidence in *E. coli*) in *S. symbiotica*. Domains in each frame are represented by coloured boxes, with similar colours used for repeated domains in each protein. on top of each box, the domain’s is provided. Above each frame, the protein’s name is stated.On the bottom-right of every frame, a scale bar is provided.

## Conclusion

*S. symbiotica* strains analysed here have all been evolving under a similar environment, the aphid-*Buchnera* symbiotic system. We have established that *S. symbiotica* strains can be considered to be along the genome reduction spectrum from a free-living bacterium to a drastically-reduced endosymbiont, thus providing “snapshots” of the genome reduction process. SAf would thus represent the very first stages of genome reduction, having not yet lost its ability to be grown in axenic culture and having undergone a mild genome shrinkage and few rearrangements, when compared to the free-living Db11. SAp and SCt would be a stage further down the path, having a more reduced genome than SAf and showing a massive enrichment in both pseudogenes and MEs. However, SCt has already done the transition to becoming a co-obligate endosymbiont, and thus shows more drastic gene losses in the EAAs’ biosynthetic pathways. SCc and STs find themselves in more advanced stages of genome reduction and integration to their symbiotic systems, having established a series of metabolic dependencies and complementation with *Buchnera* for the synthesis of several essential compounds. Nonetheless, SCc differs greatly from STs in genome size, which is explained by the former being in a recent stage of an advanced genome erosion, thus retaining several pseudogenes and “junk” DNA. Both SCc and STs display a drastic genome-wide gene loss, and particularly in their ncRNA repertoire and informational machinery. Through the comparison of these *S. symbiotica* strains, we were able to hint at essential retained functions, which not surprisingly are shared with other highly-reduced endosymbionts. The detailed study of protein diminution in *S. symbiotica* revealed a common tendency of endosymbionts to loose non-essential protein domains, and thus constituting an additional route towards genome reduction. We expect the further study of this particular endosymbiont of aphids will continue to provide important clues into the intriguing process of genome reduction.

## Methods

### Annotation of protein-coding genes

All protein-coding genes that were not found in their respective *S. symbiotica* genomes were searched for using the online version of **tblastn**^74^ with *S. marcescens*’ or *E. coli*’s protein as query. All positive hits with an e-value ≤10^-3^ were then manually curated. Domains within a protein were annotated using the **InterProScan**^75^ webserver and through alignments against *E. coli*’s proteins using **MAFFT** v7.220^76^. Circular representations of presence/absence of genes were done using **circos** v0.67^77^ and edited in Inkscape v0.91. COG categories for proteins were assigned using **blastx** and *ad hoc* perl scripts to select the best non-overlapping hits with an e-value threshold of ≤10^-3^. Then, COG categories absolute counts were converted to relative ones per organism. Finally, in order to analyse the disturbance of *S. symbiotica*’s functional profiles from that of free-living Db11, we subtracted Db11’s relative frequency per COG from each value within the same row. Visual display of COG categories was done using **R** and the **gplots** library, followed by manual editing in Inkscape.

### Annotation of RNA features

tRNA features were annotated using **tRNAscan-SE** v1.3.1^78^ (-B option for the bacterial model) and **TFAM** v1.4, followed by manual curation. All other RNA features were searched for using **Infernal** v1.1.1^79^ (--cut_tc --mid) against the **Rfam** v12.0 database^80^. All hits with an e-value lower than -10^-3^ were considered and manually curated. Visual displays were done using **R** and the **gplots** library, followed by manual editing in Inkscape. Plain-text source files used for the plotting of RNA features can be found in https://figshare.com/s/193f7513676a83b52241.

### Rearrangement analysis

Single copy shared proteins among *S. symbiotica* strains and Db11 were calculated as in^37^. Briefly, we used **OrthoMCL** v2.0.9^81^ to build the orthologous groups of proteins, followed by manual curation aimed at joining rapidly-evolving proteins such as outer membrane proteins. This proteins were then used as rearrangement markers for calculating a minimal rearrangement phylogeny in **MGR** v2.0.3^82^ (no heuristics). Scaffold for unfinished genomes (SAf, SAp, and SCt) were arranged as to minimise the distance against Db11. Tree visualisation was done in **FigTree** v1.4.1. Rearrangement graphic was done in **R** using the **genoPlotR** library^83^. All graphics were edited in Inkscape.

## Acknowledgements

This work has been funded by the Ministerio de Economia y Competitividad (Spain) co-financed by FEDER funds [BFU2015-64322-C2-1-R to A.L.]; the European Commission [Marie Curie FP7 PITN-GA-2010-264774-SYMBIOMICS to A.M.M]; and the Consejo Nacional de Ciencia y Tecnologia (Mexico) [Doctoral scholarship CONACYT 327211/381508 to A.M.M.]. The funders had no role in study design, data collection and analysis, decision to publish, or preparation of the manuscript.

## Author contributions statement

A.M.M. and A.L. conceived and designed the study. A.M.M. analysed the data. A.M.M. and A.L. wrote, reviewed, and approved the final version of the manuscript.

## Additional information

### Accession codes

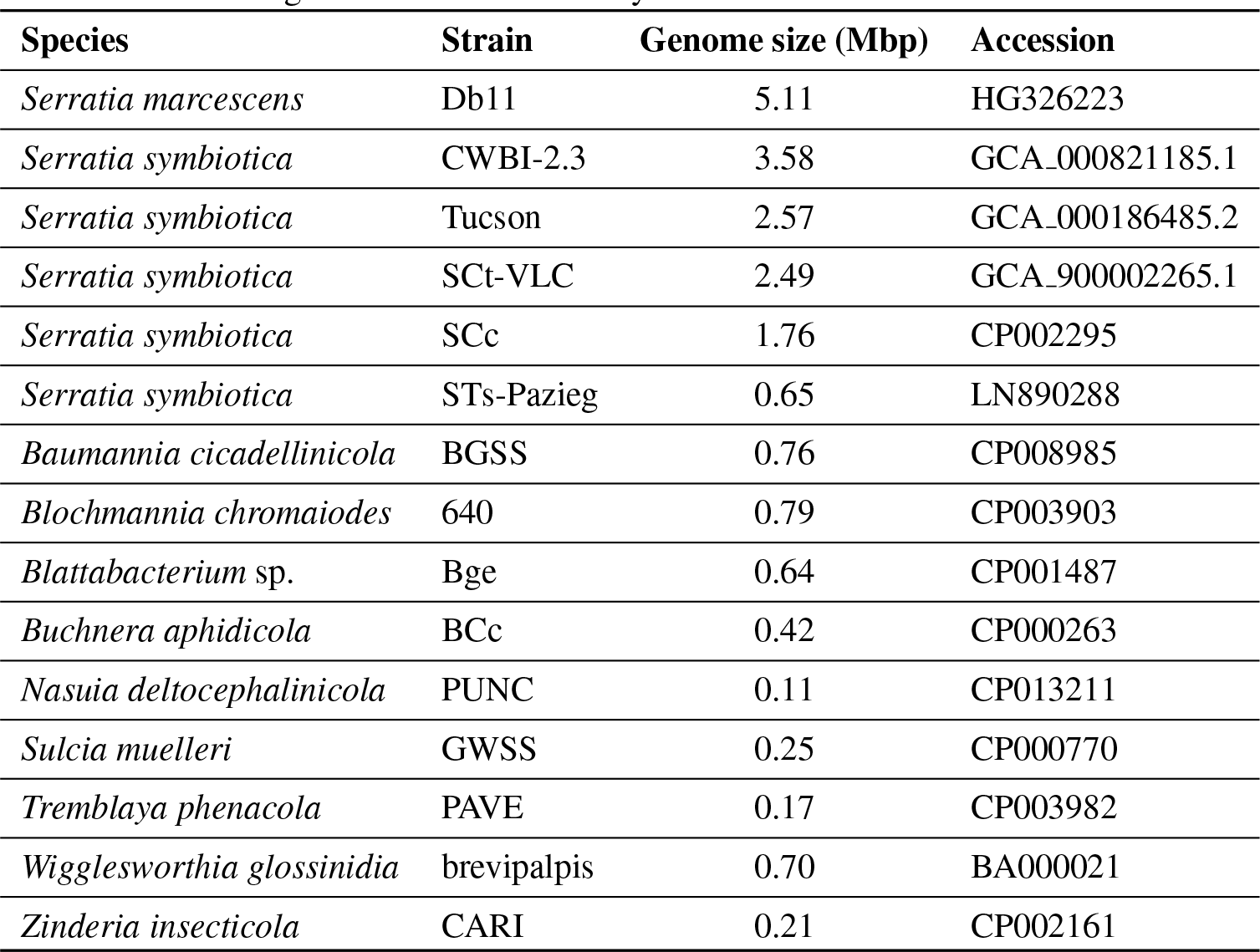
Accessions for all organisms used in this study.

| Species | Strain | Genome size (Mbp) | Accession |
| --- | --- | --- | --- |
| *Serratia marcescens* | Db11 | 5.11 | HG326223 |
| *Serratia symbiotica* | CWBI-2.3 | 3.58 | GCA_000821185.1 |
| *Serratia symbiotica* | Tucson | 2.57 | GCA_000186485.2 |
| *Serratia symbiotica* | SCt-VLC | 2.49 | GCA_900002265.1 |
| *Serratia symbiotica* | SCc | 1.76 | CP002295 |
| *Serratia symbiotica* | STs-Pazieg | 0.65 | LN890288 |
| *Baumannia cicadellinicola* | BGSS | 0.76 | CP008985 |
| *Blochmannia chromaiodes* | 640 | 0.79 | CP003903 |
| *Blattabacterium* sp. | Bge | 0.64 | CP001487 |
| *Buchnera aphidicola* | BCc | 0.42 | CP000263 |
| *Nasuia deltocephalinicola* | PUNC | 0.11 | CP013211 |
| *Sulcia muelleri* | GWSS | 0.25 | CP000770 |
| *Tremblaya phenacola* | PAVE | 0.17 | CP003982 |
| *Wigglesworthia glossinidia* | brevipalpis | 0.70 | BA000021 |
| *Zinderia insecticola* | CARI | 0.21 | CP002161 |

## Competing financial interests

The authors declare no competing financial interests.

